# Syndecan functions to regulate Wnt-dependent axon guidance in *C. elegans*

**DOI:** 10.1101/046094

**Authors:** Samantha N. Hartin, Meagan Kurland, Brian D. Ackley

## Abstract

Cell adhesion molecules are key to axon guidance during development, for example specific cues can instruct axons to terminate in a specific area, or to continue growth. Syndecans are conserved cell-surface receptors that function in multiple developmental contexts. *Caenorhabditis elegans* with mutations in the single syndecan gene, *sdn-1,* exhibited errors in anterior-posterior guidance, with axons that stopped short of, or grew past their stereotypical termination point. Syndecan function was cell non-autonomous for GABAergic axon outgrowth during early development, but was likely cell autonomous to inhibit growth later in development. *sdn-1* appeared to regulate the inhibitory activity of the *egl-20/Wnt* ligand. Removing *egl-20* from *sdn-1* mutants resulted in fewer animals with prematurely terminating axons. The proteoglycan modifying enzymes *hse-5* and *hst-2*, but not *hst-6*, had similar effects, suggesting specific heparan sulfate modifications regulated EGL-20 axon-terminating activity. *sdn-1* functioned with *lin-17/Frizzled*, *bar-1*/β-catenin and the *egl-5* Hox-like transcription factor in EGL-20-depedent axon outgrowth. *bar-1* was required for *egl-5* expression in the most posterior GABAergic neurons. *sdn-1* mutations did not eliminate *egl-5* expression, but over-expression of *egl-5* rescued *sdn-1* phenotypes. Our results suggest syndecan is a component of Wnt-signaling events that are necessary for axons to recognize appropriate termination points.

## Introduction

Nervous system development is an ordered and complex process that relies on axons interpreting multiple molecular cues to form functional circuits. Cues in the extracellular matrix (ECM) are essential to neurons forming appropriate networks. For example, ephrins and their receptors, EphRTKs, are key mediators of precise axon pathfinding in retino-tectal mapping (Drescher *et al.* 1997; Thakar *et al.* 2011). Also, some guidance cues, for example Netrins, can serve as both attractants and repellants, depending on other extracellular factors, or the receptors present on the cell surface (Kennedy *et al.* 1994; Colamarino and Tessier-Lavigne 1995; Chan *et al.* 1996; Wadsworth and Hedgecock 1996). Our complete understanding of how guidance cues orchestrate neural development remains incomplete. To that end, *Caenorhabditis elegans* provides a valuable model to study nervous system development. Growth and termination of the GABAergic motorneurons have identified novel axon guidance factors and/or new roles for known guidance cues (Wightman *et al.* 1997; Hobert *et al.* 1999; Huang *et al.* 2002; Huang *et al.* 2003; Huarcaya Najarro and Ackley 2013; Opperman and Grill 2014).

Heparan sulfate proteoglycans and chondroitin sulfate proteoglycans (HSPGs and CSPGs, respectively) are crucial for proper axon guidance (Yamaguchi 2001; Lee and Chien 2004; Rawson *et al.* 2005; Rhiner *et al.* 2005; Bulow *et al.* 2008; Diaz-Balzac *et al.* 2014; Shen 2014). HS- and CSPG side chains are enzymatically modified to alter function (Bulow and Hobert 2004; Wang *et al.* 2015). For example, modified HS- or CS-side chains regulate the binding affinity and avidity for multiple signaling molecules including Wnts, FGFs and BMPs. As a functional consequence, HSPGs and CSPGs can create growth factor reservoirs in the ECM or function as co-receptors on the cell surface (Baeg and Perrimon 2000; Tumova *et al.* 2000; Yamaguchi 2001; Aricescu *et al.* 2002; Inatani *et al.* 2003; Steigemann *et al.* 2004; Bishop *et al.* 2007).

Syndecans are a major family of HSPGs. Most mammalian genomes encode for four syndecan proteins, while the *C. elegans* genome encodes one, named *sdn-1*. SDN-1 functions autonomously in the <u>H</u>ermaphrodite <u>S</u>pecific <u>N</u>eurons (HSNs) to control cell migration and axon outgrowth and in <u>D</u>orsal <u>D</u>-type and <u>V</u>entral <u>D</u>-type motorneurons (DDs and VDs, respectively) to guide commissure growth along the dorsal-ventral axis (Rhiner *et al.* 2005). SDN-1 regulates the guidance of non-neural tissues including the distal tip cells (DTC) that migrate along the anterior-posterior axis to shape the gonads during development.

Syndecan has been implicated in Wnt signaling in vertebrates and invertebrates alike (Alexander *et al.* 2000; Schwabiuk *et al.* 2009; Dejima *et al.* 2014). In *C. elegans* DTC guidance by SDN-1 interacts with the Wnt ligand, EGL-20. In early embryogenesis SDN-1 functions in a MOM-2/Wnt-dependent event to direct cell specification (Dejima *et al.* 2014). We have previously demonstrated that SDN-1 and LIN-44 function in genetically-redundant pathways during gastrulation (Hartin *et al.* 2015). Together, these indicate SDN-1 functions in multiple Wnt growth factor signaling events. Here we expand those results by analyzing the interaction of SDN-1 and Wnt signaling in the control of axon termination.

## Materials and Methods

### Strains and Genetics

N2 (var. Bristol) was used as the wild-type reference strain in all experiments. Strains were maintained at 18-22 °C, using standard maintenance techniques as described (Brenner 1974). Alleles used in this report include: *LGI, lin-44(n1792), lin-17(n671); LGII, mig-5(rh94); LGIII, egl-5(n945); LGIV, egl-20(lq42), egl-20(gk453010); LGV, mom-2(or77); LGX bar-1(ga80), sdn-1(ok449), sdn-1(zh20), hse-5(tm472), hse-5(lq49), hst-2(ok595), hst-6(ok273).* The following integrated transgenic lines were used: *lhIs47* [*Punc-25::mCherry*]*, juIs76* [*Punc-25::gfp*]*, oxIs12 [Punc-47::gfp], muIs32* [*Pmec-7::gfp*], *juSi119* [*Psdn-1::SDN-1cDNA::GFP::sdn-1 3’UTR*] (Dejima *et al.* 2014), *wpSi12* [*Psdn-1::SDN-1::sdn-1 3’UTR*] (Edwards and Hammarlund 2014), *lqIs80* [*Pscm::gfp*], *wpIs54* [*Pegl-5::EGL-5::GFP-3xFLAG*] (Niu *et al.* 2011), and *lhIs93 [Punc-25::egl-5]*. The following transgenic extrachromosomal arrays were used: *lhEx435* [*Punc-25::mCherry* - 5 ng/μl]; *lhEx457* [*Plin-17::lin-17::rfp* - 2 ng/μl]; *lhEx350, lhEx353,* [*Pegl-20::egl-20::gfp* - 5 ng/μl]; *lhEx495* [*Punc-25::sdn-1* - 1 ng/μl]; *lhEx492, lhEx493* [*Punc-25::sdn-1* - 5 ng/μl]; *lhEx521, lhEx522* [*Punc-25::sdn-1::gfp* - 1 ng/μl], *lhEx548, lhEx549* [*Punc-25::sdn-1(ΔEFYA)::gfp* – 1 ng/μl] and *lhEx601, lhEx602* [*Punc-25::mCherry::mig-5::unc-43 3 ‘UTR –* 2 ng/μl].

### Plasmid Construction

A *sdn-1* cDNA was obtained from a cDNA library, initially transcribed by random hexamers and Superscript III (Life Technologies). The primers used to amplify the cDNA were as follows: *sdn-1cDNAF1* 5’ – caggtgattacaccaacaagac – 3’ and *sdn-1cDNA R1* 5’- cagataagtgccatcagaaacc – 3’. The resulting PCR product was cloned into the pCR8/GW/Topo entry vector (Life Technologies) to make pEVL424, and then recombined into the *Punc-25* destination vector (pBA153) using LR recombinase (Life Technologies) per the manufacterer’s protocol to generate pEVL449 (*Punc-25::sdn-1*). To generate pEVL454 (P*unc-25::sdn-1::gfp*) we cut pEVL424 with *BstBI,* and ligated with the gfp cassette from pPD113.37 (a gift of A. Fire) with *Cla*I. Any additional cloning details are fully available upon request. The *egl-5* genomic locus was amplified from wild-type genomic DNA using the following primers: *egl-5F1* 5’ – ttggaaagcagtgagagtgag – 3’ and *egl-5R1* 5’ – ggagggatcattgagaaacttgag – 3’. The PCR product was TOPO cloned intp pCR8/GW/TOPO to create pEVL483 and recombined into pBA153 by L/R clonase to make pEVL479 (*Punc-25::egl-5*). A *mig-5* cDNA was amplified from a random cDNA library, using the following primers 5’ – <u>caagcaccagtatctgcatt</u>atggagccgccatgca – 3’ and 5’ – <u>agaaggttcttctagccctt</u>cgcgttctgctgttg – 3’ and inserted via the NEBuilder (NEB) enzymatic mixture into pEVL387 [Punc-25::mCherry::unc-43 3’UTR], digested with *Xba*I. The underlined sequences above supply the overlap with the pEVL387 vector.

### Fluorescence microscopy

Axon termination (GABAergic neurons) was visualized using *juIs76, oxIs12, lhIs47* or *lhEx435* or (mechanosensory neurons) *muIs32*. Scoring and imaging were done using an Olympus FV1000 laser-scanning confocal microscope with the Fluoview software. Animals where the DNC bundle did not grow to the posterior edge the cell bodies located on the VNC, were considered under-extended. Animals in which the DNC axons grew past the cell bodies located on the VNC, were scored as over-extended. In cases where the VD motorneuron cell bodies were displaced the position of the anus was used as a secondary landmark. Animals with posterior neurites (Pdns) or other obvious axon guidance errors during commissural growth were not included in the analysis of DNC termination points.

### Statistics

Fisher’s exact test was used to evaluate the statistical significance between genotype pairs. P values were calculated with Prism GraphPad (5.0) program or the GraphPad QuickCalcs (http://www.graphpad.com/quickcalcs/). A multiple test correction (Bonferroni) was applied when used to evaluate relationships within groups of genotypes, relative to the number of comparisons being made within that experimental design.

### Data Availability

All strains and plasmids presented are available upon request.

## Results

### Syndecan is an anterior/posterior axon guidance factor

The GABAergic D-type motorneurons cell bodies are located along the ventral nerve cord (VNC). The six Dorsal D-type (DD) and thirteen Ventral D-type (VD) cell bodies extend a single neuronal process anteriorly. The axon stalls, and initiates a dorsal commissural process and then a second process initiates from the stall point and extends a second anterior branch to the next most anterior VD neuron (Figure 1) (Huarcaya Najarro and Ackley 2013). The commissural process, upon reaching the dorsal nerve cord (DNC), bifurcates and sends processes both anteriorly and posteriorly. In the VNC and DNC, the VD processes fasciculate with those of the DD neurons and terminate at stereotyped positions along the anteroposterior axis (White *et al.* 1986). The most posterior DD and VD processes extend as a bundle that terminates at the anteroposterior position superior to the locations of the DD6 and VD13 cell bodies, which are located on the ventral side (Maro *et al.* 2009) (Figure 1A and B). The stereotyped development allows for detection of mutants with aberrant axon patterning phenotypes.

**Figure 1.**
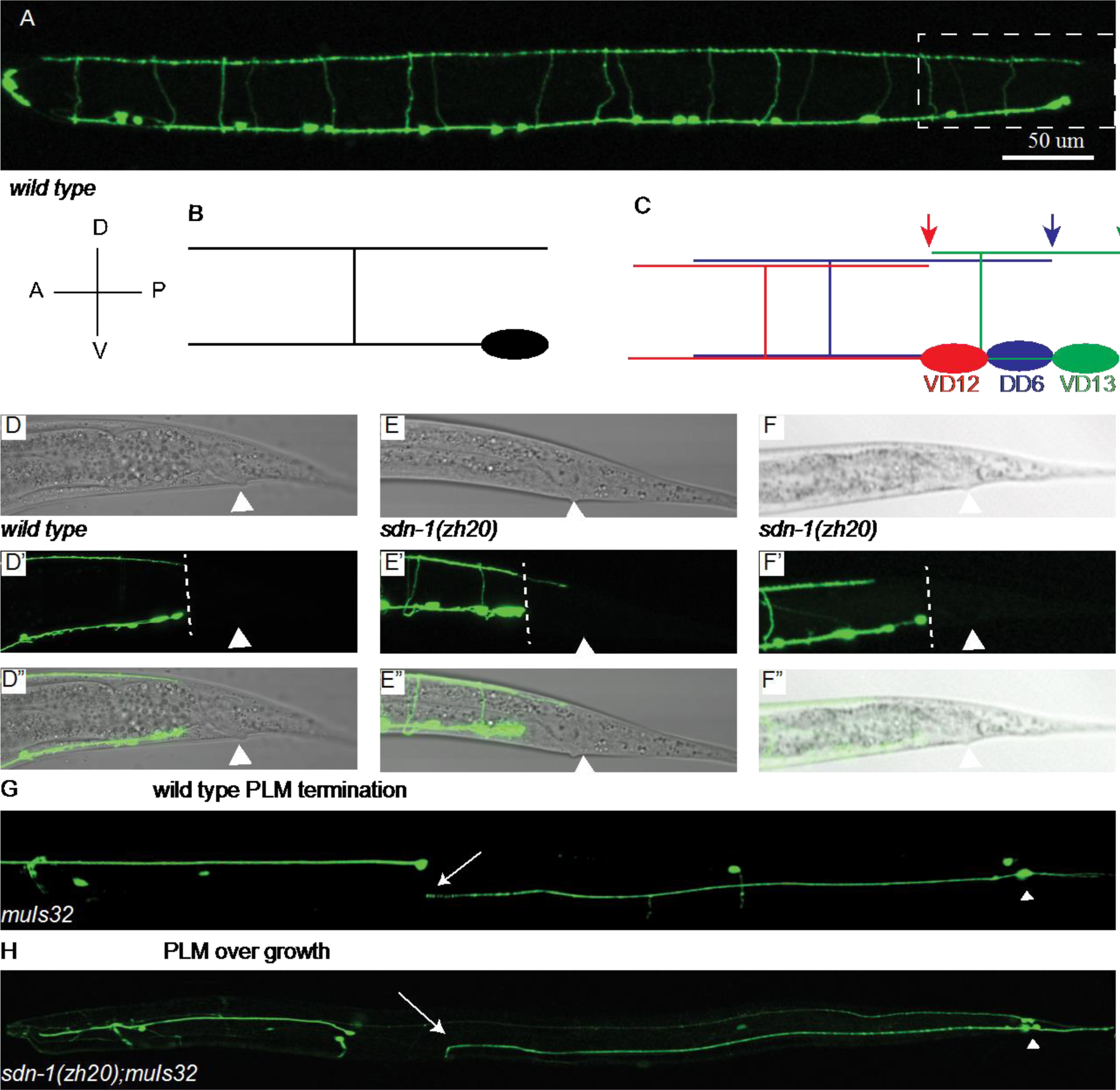
*sdn-1* regulates VD13 termination in the DNC. A) In an L4 wild-type *juIs76* [*Punc-25::gfp*] animal it is possible to visualize the pattern of axon growth of the GABAergic motorneurons, anterior is left and ventral down in all images. Boxed region in (A) is the tail region equivalent to those presented in (D-F). B) A schematic of the expected morphology of a generic D-type or V-type GABAergic motorneuron, the posterior branch of the dorsal projection terminates approximately at the posterior edge of the cell body. C) A schematic of the terminal three neurons, VD12, DD6 and VD13 and approximate termination points are indicated by arrows. D) A wild type L4 animal where the dorsal nerve cord terminates in a line with the cell bodies (dashed line). The arrowhead indicates the position of the anus. E) An *sdn-1(zh20)* L4 animal with an over-extended DNC, progressed past the plane of the last cell body, and is approximately at the position of the anus. F) An *sdn-1(zh20)* L4 animal with an under-extended DNC, terminating at approximately the position of the DD6 cell body. In wild-type *muIs32* animals the PLM (arrowhead indicates PLM cell body) axons terminate (arrow) along the anterior-posterior axis around the position of the ALM soma. B) In *sdn-1(zh20)* mutants we find that some PLM axons reach the normal termination point and then turn ventrally to continue growth, over-extending. C) Loss of function in *lin-44*/Wnt can result in PLM axons that grow posteriorly and then turn anteriorly, making the PLM appear reversed. D) *lin-44* mutants can also display PLMs were the anterior branch and posterior branch are of approximately equal length. E) The number of animals presenting with each type of PLM growth defect.

We observed that *sdn-1(zh20)* mutants had defects in formation of the GABAergic motorneurons along the anterior-posterior (A-P) axis of the animal. Animals with mutations in *sdn-1* exhibited posteriorly-directed neurites (Pdns) (16%±7), where an axonal process was inappropriately projected toward the posterior of the animal (Huarcaya Najarro and Ackley, 2013). In animals where the axon projected normally toward the anterior we also observed defects where axons terminated along the A-P axis in the dorsal nerve cord (Figure 1). We found that 35%±13 of *sdn-1(zh20)* mutants had axons that stopped short of the stereotyped termination point, while 26%±10 had axons that extended posteriorly past the proper termination point (Figure 1 D-F, Table 1). It is worth noting that in other genetic contexts where D-axon termination has been investigated it is common to observe both under- and over-extension defects in the same genotypes (Maro *et al.* 2009; Opperman and Grill 2014).

**Table 1.** GABAergic DNC termination defects in Syndecan mutants

| | % Observed (Mean ± St. Dev) | | | | Comparison to *sdn-1(zh20)* | |
| --- | --- | --- | --- | --- | --- | --- |
|  | Under | WT | Over | N | P (under) | P (over) |
| *sdn-1(zh20)* | 35 ± 13 | 39 ± 10 | 26 ± 10 | 246 | | |
| *sdn-1; juSi119* | 6 ± 1 | 36 ± 3 | 59 ± 4 | 200 | <0.0001 ‡ | <0.0001 ‡ |
| *sdn-1; wpSi12* | 1 ± 1 | 61 ± 8 | 38 ± 9 | 216 | <0.0001 ‡ | 0.0538 |
| *sdn-1; Punc-25::sdn-1 (A)* | 37 ± 13 | 52 ± 8 | 11 ± 6 | 81 | 0.6038 | 0.0035 ‡ |
| *sdn-1; Punc-25::sdn-1 (B)* | 33 ± 3 | 57 ± 6 | 10 ± 3 | 141 | 1 | 0.005 |
| *sdn-1; Punc-25::sdn-1::gfp* | 33 ± 4 | 67 ± 4 | 0 ± 0 | 104 | 1 | <0.0001 ‡ |
| *sdn-1; Punc-25::sdn-1(ΔEFYA)::gfp* | 30 ± 6 | 43 ± 1 | 28 ± 5 | 159 | 0.3844 | 0.4155 |

‡ Significantly different ( $P < 0.0042$ )

We also tested a second LOF allele, *sdn-1(ok449)* which results in an in-frame deletion that produces a shortened form of SDN-1. This shortened form lacks two of the major conserved heparan sulfate attachment sites in the extracellular domain (Minniti *et al.* 2004). We found that animals with the *ok449* mutation had similar defects to those found with the *zh20* allele (Table 1). We continued our studies using the *zh20* allele since it has been demonstrated to be a genetic and molecular null (Rhiner et al., 2005, Dejima et al., 2014).

The VD neurons are formed at the end of the first larval stage (L1), and finish their initial outgrowth during the beginning of the second larval stage (L2). We initially scored animals in the fourth larval stage (L4). Between the L2 and L4 stages the animals approximately double in length. During development, the posterior segment of the DNC (from the point of the final commissural branch to the terminus) increases from ~25-30 μm (L2) to ~30-40 μm (L4), and continues growing to ~50-60 μm in wild-type young adults. To determine if *sdn-1* axon outgrowth errors were due to developmental errors or a failure to keep up with the growth of the animal we scored DNC termination in *zh20* animals at the L2 stage. We found that like those scored at later stages, *zh20* animals at the L2 stage had both under-(52%) and over-(27%) extended GABAergic axons in the DNC. We concluded from this analysis that, during initial formation, axons terminated growth aberrantly in most animals lacking *sdn-1*.

To determine if the effects of axon termination along the anterior-posterior axis were limited to the D-type neurons we examined the axons of the six mechanosensory neurons using an integrated GFP reporter, *muIs32.* In wild-type animals the <u>A</u>nterior <u>L</u>ateral <u>M</u>echanosensory (ALMs) and the <u>P</u>osterior <u>L</u>ateral <u>M</u>echanosensory (PLMs) neurons extend anteriorly-directed axons from their cell bodies and have a characteristic termination point, with the PLMs terminating just posterior to the ALM cell bodies (Figure 1) and the ALMs terminating just posterior to the nose of the animal. In addition, ALM and PLM axons form largely parallel to the dorsal and/or ventral nerve cords, and do not appear to deviate in the dorsal-ventral axis, except where they make branches into the ventral nerve cord (PLMs) or the nerve ring (ALMs).

When we examined the ALM and PLM neurons in *sdn-1(zh20)* mutant animals we found disruptions in the outgrowth of the neurons. As had been previously reported the ALM cell bodies were frequently displaced posteriorly, likely due to incomplete cell migration (Rhiner *et al.* 2005). Because of migration defects, we did not score axon termination errors in the ALMs. The PLM neurons do not undergo long range cell migrations, and were in grossly normal positions in *zh20* mutants. We found PLM axons that were shortened in *zh20* animals (14%±4, N=207), either by terminating prematurely or not elongating sufficiently during organismal growth. We also found instances where PLMs were over-extended, and could “hook” toward the ventral nerve cord (17%±6) (Figure 1) (Grill *et al.* 2007). Overall these defects were less penetrant than those we observed in the DD/VD motorneurons, but were consistent with a role for *sdn-1* in regulating axon extension along the A/P axis, and that loss of *sdn-1* can result in errors in the precision with which axons terminate in different neuron types.

### Syndecan function is required in multiple tissues to regulate D-type neuron outgrowth

We partially rescued the outgrowth phenotypes in the D-type neurons in *sdn-1(zh20)* animals by reintroducing *sdn-1* (Table 1
). We obtained a Mos1-mediated single copy insertion (MosSCI), *juSi119,* in which a GFP-tagged version of SDN-1, under the regulation of *sdn-1* endogenous promoter elements, had been inserted on the second chromosome (LG II) (Dejima *et al.* 2014). The *zh20; juSi119* animals had fewer undergrown axons in the DNC (Table 1), but had a higher rate of DNC overgrowth (59%±4) than was observed in *zh20* mutants. To determine if this was due to a detrimental effect of the GFP insertion we obtained a second MosSCI, *wpSi12*, which is inserted in the same chromosomal position, but is not GFP tagged (Edwards and Hammarlund 2014). We again observed rescue of the under-extension phenotype, but increased overgrowth (Table 1).

To test whether *sdn-1* could be functioning cell-autonomously we drove expression of *sdn-1* specifically in the GABAergic neurons using the *unc-25* promoter (1 ng/μl). Cell-specifically expressed *sdn-1* partially rescued the over-extension phenotype in *zh20* mutants in two separate lines (26% - *zh20* vs. 11%, P=0.0035 and 10%, P=0.005). In both transgenic lines the rate of under-extended axons was not significantly changed (35% - *zh20* vs. 37%, P=0.60 and 33%, P=0.91). No obvious differences were found in lines generated at a higher concentration (5 ng/μl). We were still unable to rescue the under-extension defects (36% vs 34%, P=0.94) and there was not a robust increase in the efficiency of rescuing the over-extension defects (11% vs. 8%, P>0.05). We also cell-specifically expressed an SDN-1::GFP chimera in the GABAergic motorneurons, and found that we could again rescue over-extension, but not under-extension. These data suggest that the over-extension of axons in *zh20* animals was due to, at least in part, the cell-autonomous loss of *sdn-1*, but the under-extension phenotype demonstrated a requirement for *sdn-1* function in other tissues.

Syndecans have been shown to be linked to cytoplasmic effectors via their PDZ binding motif. To determine if the conserved, potential PDZ binding motif (EFYA) at the very C-terminus was required for SDN-1 function in the D-neurons, we deleted it from our SDN-1::GFP. We cell-specifically expressed the SDN-1::GFP(ΔEFYA) in the GABAergic motorneurons, and found that, unlike the receptor with the intact EFYA motif, this version failed to rescue the over-extension phenotype in *sdn-1(zh20)* animals (Table 1). This suggests that, intracellularly, SDN-1 may link to effectors that are important for axon termination via the EFYA motif.

### Syndecan is enriched in the DNC near the VD13 termination point

We examined the localization of SDN-1GFP expressed by *juSi119.* The protein was present broadly throughout the animal. We observed robust expression in the DNC and tissues surrounding it, including the muscle, epidermis and intestinal cells (Figure 2). We further examined whether SDN-1 was expressed in the D-type neurons using a cytoplasmic RFP (*Punc-25::mCherry*). We found that, while SDN-1GFP was present in the ventral and dorsal nerve cords, we were unable to detect robust expression in the D-type neurons. We examined animals specifically when the VD13 commissure was being formed, and found no detectable SDN-1::GFP in the growth cone (Figure 2). This was curious since previous work has suggested the *sdn-1* promoter is active in the D-type neurons (Rhiner et al., 2005). Since we were unable to rescue the overgrowth phenotype with the MosSCI integrated transgenes, but we could with cell-autonomous expression, we inferred the MosSCI integrated SDN-1 transgenes into LGII may have been poorly expressed in the GABAergic neurons.

**Figure 2.**
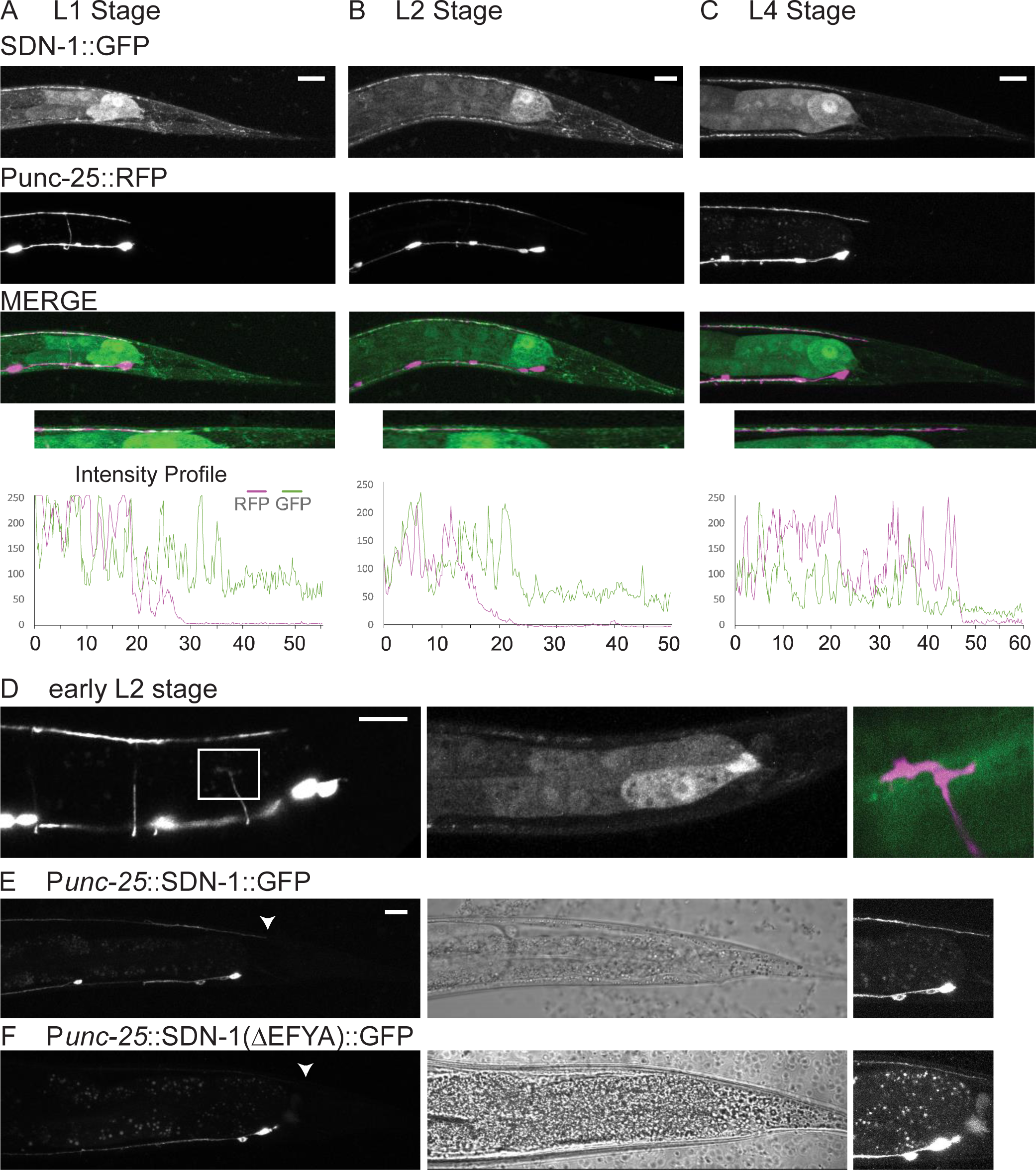
SDN-1::GFP accumulates near the D-type termination point in the dorsal nerve cord. SDN-1::GFP expression (*juSi119*) relative to the D-type neurons (*Punc-25*::RFP). A) In wild-type first larval stage animals (L1) GFP was present in the tail in multiple tissues, and was localized to the dorsal nerve cord. Line scans were generated by drawing a 1-pixel wide line along the dorsal nerve cord, and measuring the intensity using the Plot Profile command in ImageJ. The graphs illustrate the accumulation of SDN-1 near the end of the GABAergic DNC. No gross differences in the accumulation of SDN-1 were observed as the animals aged through the L2 (B) and L4 (C) larval stages. D) SDN-1::GFP was not obviously localized to the growth cone of the VD13 neuron during outgrowth. E) Expression of an SDN-1::GFP chimera specifically in the D-type neurons. SDN-1 localized primarily to the dorsal and ventral nerve cords, and was present near the expected termination (arrowhead). F) Removal of the EFYA motif from the SDN-1::GFP resulted in a reduced localization to the dorsal cord. At the right, the contrast enhanced image demonstrates that, although reduced, the SDN-1(ΔEFYA)::GFP was able to localize to the dorsal cord.

While we did not detect abundant SDN-1::GFP in VD neurons, we did find that, in the DNC, SDN-1::GFP was enriched near the normal termination point of the GABAergic neurons (Figure 2). In L1 animals, prior to the formation of the VD13 neuron, we found that SDN-1::GFP accumulated in the nervous system, and aggregated near the termination point of the GABAergic neurons in the DNC (Figure 2). This suggests that SDN-1::GFP would be in a position to assist the later arriving VD13 axon in identifying the proper termination point. When we examined the SDN-1::GFP after all the VD axons had formed (Figure 2) we found that SDN-1::GFP was maintained at the termination point of the VD13 axon, being present in L4 (Figure 2) and adult animals (not shown).

The *Punc-25::*SDN-1::GFP transgene was capable of rescuing the dorsal cord growth defects as efficiently as the non-tagged version (Table 1). In these animals, we observed SDN-1 in cell bodies and along the ventral, commissural and dorsal processes of the neurons (Figure 2), but no specific enrichment at the termination point of the DNC. From this analysis, we concluded that SDN-1 was localized such that it could make a local contribution to GABAergic axon termination in the dorsal nerve cord, and that, within the GABAergic neurons SDN-1 could localize to the dorsal cord. We also concluded that SDN-1 accumulation near the termination point of the dorsal nerve cord observed by *juSi119* likely represented contributions from multiple tissue types.

We subsequently analyzed the localization of the SDN-1::GFP(ΔEFYA). Unlike the SDN-1::GFP, which appeared to localize evenly to the dorsal and ventral cords, the version of SDN-1 lacking the EFYA motif was biased towards accumulation in the ventral nerve cord, relative to the dorsal nerve cord (Figure 2). We cannot differentiate whether this reflects a failure in trafficking or retention of the receptor in the dorsal aspect of the neuron. However, it is consistent with the EFYA motif being required for rescue in the D-type neurons.

### SDN-1 function is epistatic to LIN-44 in VD neurons, but not PLMs

Previous work has demonstrated that Wnt signaling can regulate axon termination of the D-type neurons (Maro *et al.* 2009). Consistent with those observations we also found that animals with mutations in the most posteriorly expressed Wnt ligand, *lin-44(n1792)*, caused the DNC axons to be over-extended (Figure 3, Table 2). Previously we described that *sdn-1* and *lin-44* exhibit a synthetic lethal genetic interaction in early development, with ~75% of double mutants dying during embryogenesis (Hartin *et al.* 2015). To determine whether *sdn-1* was functioning in *lin-44* dependent outgrowth we scored the axons in the surviving animals. Unlike the *lin-44* single mutants, the doubles exhibited both under and over-extended DNCs (Figure 3, Table 2). Qualitatively, we assessed the over-extended DNCs in the *sdn-1; lin-44* mutants to be more like those of the *lin-44* animals, as they extended further into the posterior than was typically seen in *sdn-1* mutants. Thus, we concluded that *sdn-1* loss of function was partially epistatic to axon overgrowth caused by *lin-44.* We hypothesized that the undergrown axons might terminate prior to reaching the position where they would be sensitive to the presence (or absence) of *lin-44.* However, axons that did extend past the normal termination point were affected by the loss of *lin-44,* and protruded into more posterior regions of the animal.

**Figure 3.**
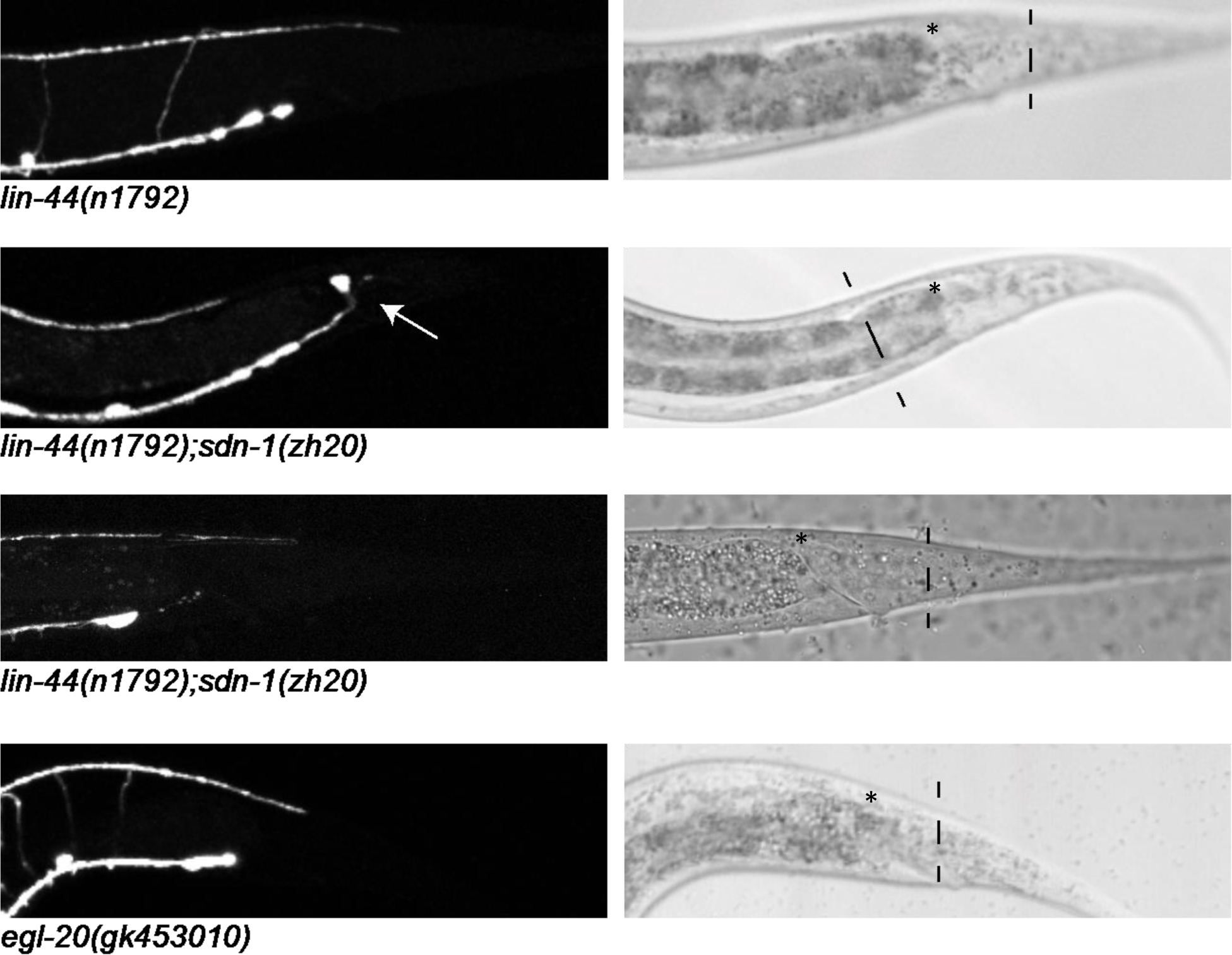
*lin-44* and *egl-20* Wnt ligands regulate the termination point of the dorsal nerve cord. Loss of function mutations in *lin-44* resulted in the DNC overextending into the posterior regions of the animal. The arrowhead indicates the expected termination point. In the DIC image the dashed line indicates the termination point of the axon fascicle in the animal. The asterisks indicate the approximate wild-type termination point. Note, the second *lin-44; zh20* animal was visualized using *oxIs12,* which is linked to *sdn-1* on LG X, and also labels the DVB neuron (arrow), which is located just posterior and slightly dorsal to the VD13 cell body. Loss of function in *egl-20* also resulted in over-extension of the DNC.

**Table 2.**
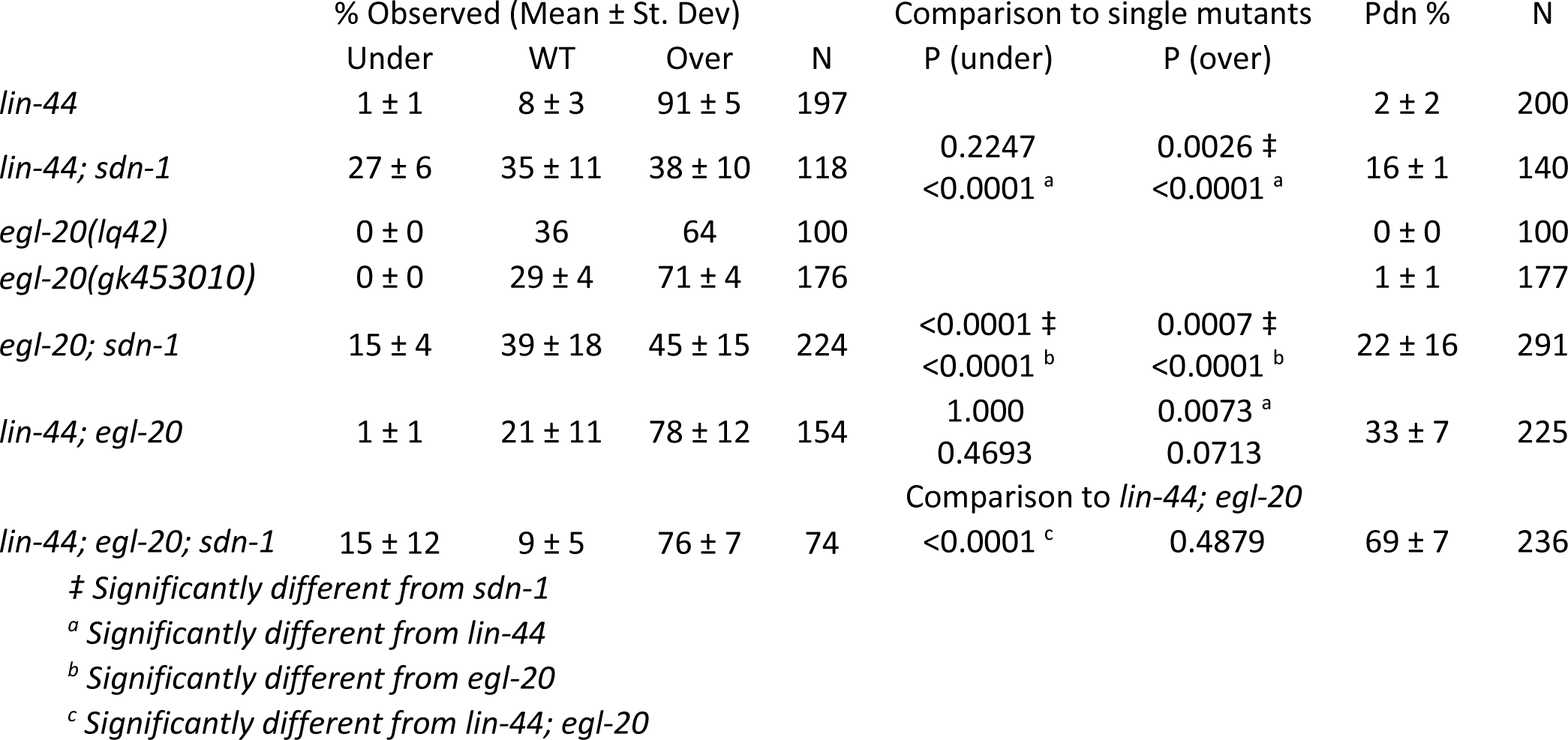
Wnt ligands and *sdn-1* DNC termination phenotypes

Mutations in *lin-44* also affect the direction and extent of axon outgrowth of the PLM axons (Hilliard and Bargmann 2006) (S. Figure 1). Loss of function in *lin-44* resulted in PLMs with posteriorly directed axons (54%±18, N=181) and altered lengths of posterior and anterior processes, including animals where the processes appeared to be of equal length (20%±15) and animals with longer anterior processes, but that stop short of the normal termination point (9%±4). The surviving *lin-44; sdn-1* animals had an increase in the number of reversed PLMs (85%±6, N=200 P<0.0001 vs *lin-44* alone). Thus, we concluded that *sdn-1* function was not epistatic to *lin-44* in PLMs, rather, in parallel. And that *sdn-1* had a similar, but weaker effect on PLM polarity as *lin-44,* but the effect was only obvious in the animals also lacking *lin-44.* To better understand how *sdn-1* functioned in axon outgrowth we focused the remainder of our study on dorsal nerve cord termination in the D-type neurons.

**Supplementary Figure 1.**
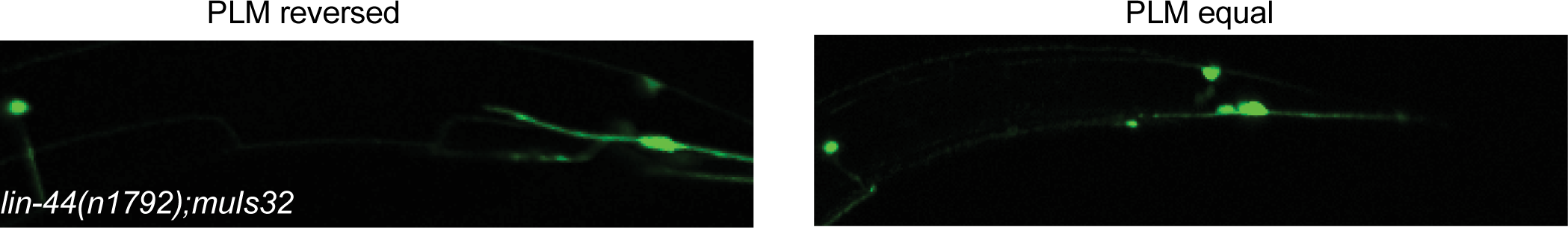
PLM defects in *lin-44* mutants. PLM neurons in *lin-44* mutants were often observed with reversed morphologies, where the posterior branch of the axon is much longer than the anterior branch. The axon projected into the tail and then turn anterior projecting past the cell body. In other animals, the anterior and posterior branches exhibit approximately equal lengths.

### EGL-20 and LIN-44 function in parallel in GABAergic development

A second Wnt ligand, *egl-20,* has also been found to affect GABAergic neuron termination in the DNC (Maro *et al.* 2009). We obtained two putative null alleles of *egl-20*, *lq42* (Josephson *et al.* 2016) and *gk453010* from the Million Mutation Project (Thompson *et al.* 2013). Both *lq42* and *gk453010* introduce premature stop codons in the *egl-20* mRNA, unlike the reference allele, *n585*, which is a missense mutation. We found that in both *egl-20(lq42)* and *egl-20(gk453010)* axons over-extended into the posterior (Table 2). We subsequently used the *gk453010* allele for future analyses. Qualitatively, we observed that the extent to which the axons extended past the expected termination point was shorter in *egl-20* than in *lin-44* mutants, which is consistent with their patterns of expression, with *egl-20* being expressed in a region just anterior to the expression of *lin-44*.

We generated *lin-44; egl-20* double mutants, and found that a higher number of animals than expected exhibited Pdn defects (27%±6, compared to 0.5% - *egl-20,* and 1.5% - *lin-44*), as well as additional guidance defects during commissural growth, making it difficult to accurately assess the DNC phenotype. In the axons that could be scored, we found that most of the animals had over-extended DNCs (Table 2). Qualitatively, as in the *sdn-1; lin-44* doubles, the length of the over-extended DNCs appeared more severe in the Wnt double mutant than in *sdn-1* single mutants, which is consistent with previous reports, using other alleles of *egl-20* (Maro *et al.* 2009). We also generated *lin-44; egl-20; sdn-1* triple mutants, which were like the *lin-44; egl-20* doubles, but exhibiting an even higher rate of Pdn defects (69±7), axon guidance errors, as well as over- and under-extended DNCs.

These results suggest there are multiple Wnt-mediated events in the development of the GABAergic neurons where EGL-20 and LIN-44 were functioning. First, EGL-20 and LIN-44 functioned in a partially redundant fashion to instruct growth anteriorly from the cell body. Subsequently, EGL-20 and LIN-44 functioned in a partially overlapping manner to regulate guidance of the axons including the where the axons would terminate in the DNC.

### SDN-1 inhibits EGL-20 axon repulsion during early DNC formation

We next examined *sdn-1; egl-20* double mutants. We found a reduction in the penetrance of the under-extension phenotype present in animals lacking both *egl-20* and *sdn-1* (Table 2). One interpretation of this was that, in *sdn-1* mutants, EGL-20 was inducing premature termination of the DNC. Previous results have found that EGL-20 diffusion is regulated by SDN-1 (Schwabiuk *et al.* 2009), and this would be consistent with EGL-20 exerting an inhibitory effect with a longer range of action when SDN-1 was absent. We tested this by expressing an EGL-20::GFP from its endogenous promoter. We found that D-type axons terminated near the site of EGL-20::GFP accumulation (Figure 4). Further, we found that over-expression of EGL-20 induced mild under-extension phenotypes in otherwise wild-type animals (16%±7). We crossed the transgenes into animals lacking *sdn-1.* The penetrance of under-extension defects increased (59%±11), which was significantly higher than expected for animals simply lacking *sdn-1* (P<0.0001). These results indicate that over-expression of EGL-20 could induce premature termination of the GABAergic neurons in a syndecan-dependent manner.

**Figure 4.**
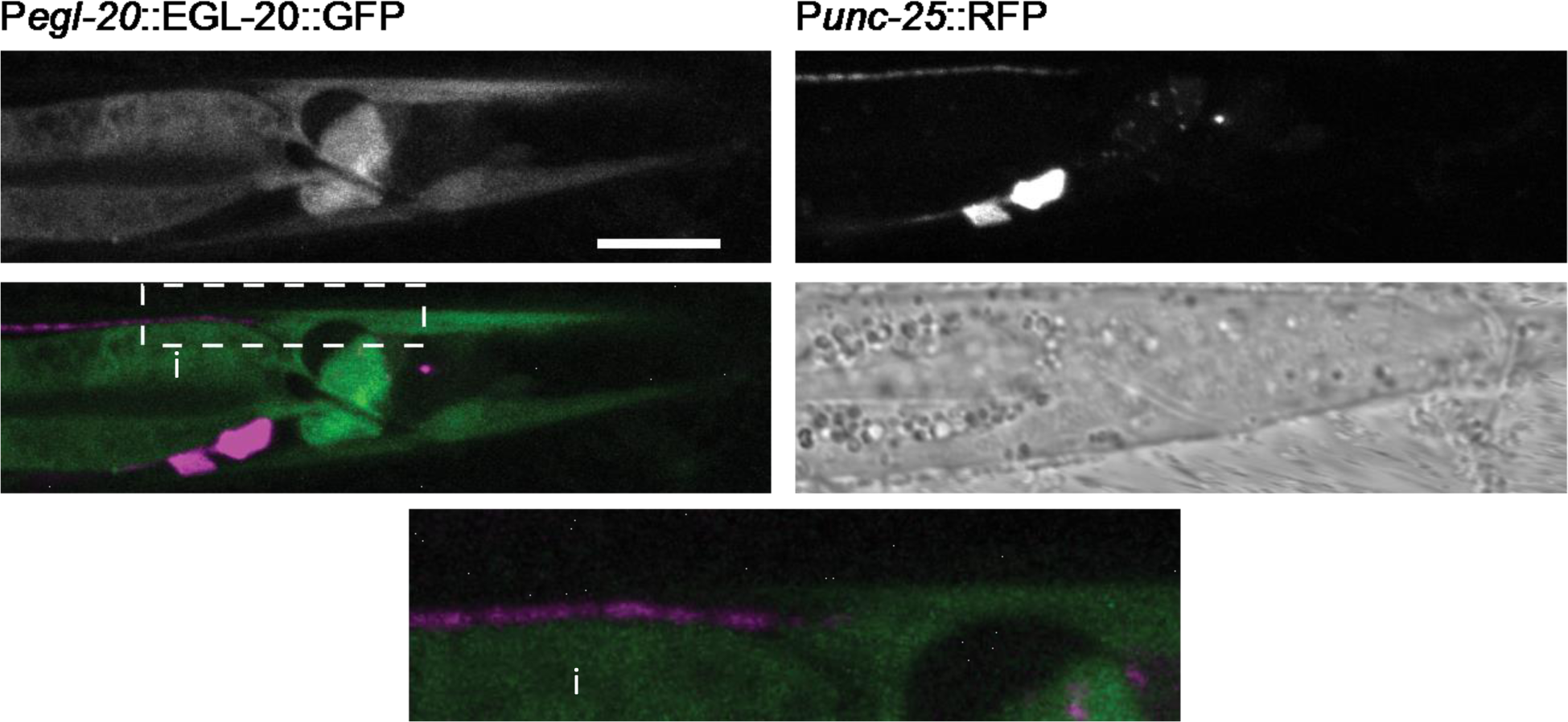
The GABAergic neurons terminate near regions where EGL-20 accumulates. EGL-20::GFP localized to the posterior of the animals, near the muscles and relative to the termination point of the GABAergic neurons in L2 animals. Note the GFP in the intestine (i) is an artifact of the coinjection marker, and is intracellular.

We reasoned that, perhaps, the increased over-extension defects we observed in the integrated SDN-1 lines could also be due to the transgene interfering with normal EGL-20 activity. We reduced the amount of SDN-1 being produced by making the inserted *juSi119* transgene heterozygous. In those animals, the over-extension phenotype was reduced from 59% to 21%, suggesting the over-extension observed was SDN-1 dose-dependent.

### HS modifiers interact with LIN-44 and EGL-20 in a context-dependent manner

Interactions between Wnt ligands and HSPGs are often mediated by the side chains present on the core protein. It has been demonstrated that the side chains on SDN-1 are modified by an epimerase, HSE-5, and at least two sulfotransferases, HST-2 and HST-6 (Bernfield *et al.* 1992; Lee and Chien 2004). Each enzyme has been shown to be involved in multiple aspects of neural development through the specific modifications each makes (Bulow and Hobert 2004; Rhiner *et al.* 2005; Bulow *et al.* 2008; Diaz-Balzac *et al.* 2014). We examined the loss-of-function mutation for the three modifying enzymes to determine whether heparan sulfate sugar modifications are important for interactions with EGL-20.

HSE-5 is a C5-epimerase that is predicted to catalyze the chain-modifying epimerization of glucuronic acid to iduronic acid during heparan sulfate biosynthesis. HSE-5 is predominantly expressed in the hypodermis and intestine (Bulow and Hobert 2004). Previously, HSE-5 has been shown to function in parallel to SDN-1 for D-type motorneuron dorsally-directed commissure outgrowth and VNC fasciculation (Rhiner *et al.* 2005). Animals with loss of function in *hse-5* exhibited both under- and over-extension phenotypes in the DNC (Table 3), indicating that the activity of this enzyme does contribute to anterior/posterior outgrowth of D-type axons. Double mutants of *hse-5* and *sdn-1* had an additive effect, inducing under-extension defects in 77%±3 animals, consistent with HSE-5 having targets in addition to SDN-1.

**Table 3.** DNC phenotypes in heparan sulfate modifying enzyme genes

| | % Observed (Mean ± St. Dev) | | | N | Comparison to *HS* single mutant | |
| --- | --- | --- | --- | --- | --- | --- |
|  | Under | WT | Over |  | P (under) | P (over) |
| *hse-5* | 45 ± 2 | 29 ± 2 | 27 ± 4 | 250 | | |
| *hse-5; sdn-1* | 77 ± 3 | 17 ± 8 | 6 ± 5 | 201 | <0.0001 ‡ | <0.0001 ‡ |
| *hse-5; egl-20* | 4 ± 5 | 28 ± 4 | 69 ± 9 | 112 | <0.0001 ‡ | <0.0001 ‡ |
| *hst-2* | 17 ± 8 | 41 ± 15 | 42 ± 6 | 127 | | |
| *hst-2; egl-20* | 8 ± 12 | 34 ± 10 | 58 ± 1 | 217 | 0.2366 | 0.0251 |
| *hst-6* | 11 ± 3 | 48 ± 23 | 41 ± 20 | 137 | | |
| *hst-6; egl-20* | 2 ± 2 | 34 ± 13 | 64 ± 11 | 246 | 0.0013 ‡ | 0.0012 ‡ |

‡ Significantly different ( $P < 0.0063$ )

We subsequently asked whether loss of *hse-5* affected the penetrance of defects in EGL-20-dependent outgrowth (Table 3). The *egl-20; hse-5* double mutants had fewer under-extended DNCs, consistent with EGL-20 inducing premature termination in animals lacking HSE-5 activity. The double mutants were not significantly different for over-extended DNCs from *egl-20* single mutants, suggesting the loss of *hse-5* was not further affecting this phenotype.

HST-2 and HST-6 can add sulfate moieties to side chains after epimerization by HSE-5. During neuronal development *hst-6* is expressed in neuronal tissues and *hst-2* is expressed in the hypodermis (Bulow and Hobert 2004). We observed a lower penetrance of under-extension defects in *hst-2* (23%) and *hst-6* (13%) mutants compared to either *hse-5* or *sdn-1* mutants. However, like *hse-5* and *sdn-1,* we found that removing *egl-20* resulted in a reduction in the penetrance of under-extension defects mutants in the transferase mutants (Table 3). Overall, we concluded that loss of the modifier enzymes was similar, but not equivalent to the loss of SDN-1, consistent with other reports of the core protein and modifiers to proteoglycan functions (Dejima *et al.* 2014; Zheng *et al.* 2015b).

### *sdn-1* is not compensated by glypican-like HSPG receptors

Glypicans are a second family of membrane-associated HSPGs. In *C. elegans* there are two glypican family members, *gpn-1* and *lon-2* (Hudson *et al.* 2006; Gumienny *et al.* 2007). Recent work has found a role for *lon-2* in netrin-mediated axon guidance (Blanchette *et al.* 2015). To determine if these receptors were the proteins modified by HSE-5 to function in parallel with SDN-1 we examined loss-of-function mutations in *gpn-1* and *lon-2,* alone, and in combination with *sdn-1.* Overall, we found that removing *lon-2* alone had little effect on termination in the dorsal nerve cord. *gpn-1* mutant animals displayed a penetrant over-extension phenotype (Table 4) that was reduced when *lon-2* was removed. In contrast, in animals already lacking *sdn-1* the removal of *gpn-1* and/or *lon-2* did not increase the penetrance of the under-extension phenotype. Based on these analyses we concluded that the glypican-like receptors are not the HSE-5-modified proteins that are in parallel to SDN-1 in dorsal cord outgrowth.

**Table 4.** DNC phenotypes in heparan sulfate-decorated receptors

| | % Observed (Mean ± St. Dev) | | | N | Comparison to *sdn-1* | | % Pdn | N |
| --- | --- | --- | --- | --- | --- | --- | --- | --- |
|  | Under | WT | Over |  | P (under) | P (over) |  |  |
| *gpn-1(ok377)* | 2 ± 2 | 32 ± 8 | 66 ± 7 | 190 | <0.0001 ‡ | <0.0001 ‡ | 4 ± 7 | 202 |
| *lon-2(e678)* | 0 ± 0 | 85 ± 10 | 15 ± 10 | 107 | <0.0001 ‡ | 0.0657 | 1 ± 1 | 108 |
| *gpn-1lon-2* | 6 ± 9 | 75 ± 3 | 19 ± 9 | 84 | <0.0001 ‡ | 0.4509 | 1 ± 2 | 85 |
| *gpn-1sdn-1* | 36 ± 15 | 50 ± 6 | 16 ± 8 | 120 | 0.9068 | 0.0785 | 17 ± 10 | 142 |
| *gpn-1lon-2sdn-1* | 34 ± 5 | 50 ± 3 | 14 ± 9 | 94 | 0.8988 | 0.0529 | 27 ± 13 | 126 |

‡ Significantly different

### *sdn-1* functions in a canonical Wnt signaling pathway to promote axon extension

Wnt signaling in *C. elegans* has been well characterized. Like others systems, in a canonical signaling pathway, activation of Frizzled receptors by Wnt ligands activates the cytosolic protein Disheveled to inhibit the GSK3/Axin/APC complex, allowing β-catenin to activate the transcription of target genes with the help of TCF-LEF transcription factors (Buechling and Boutros 2011). If EGL-20 were activating the canonical signaling pathway, then we would expect loss of function in downstream components to mimic the loss of *egl-20.* Instead, consistent with previous reports, we found some Wnt pathway mutants had both under- and over-extension defects in the DNC (Table 5). Mutations in *lin-17*/Frizzled (26%±2), *dsh*-*1*/Dishevelled (28%±0) or *mig-5*/Dishevelled (55%±2) each resulted in under-extension of the DNC, similar to data previously reported (Maro *et al.* 2009).

**Table 5.**
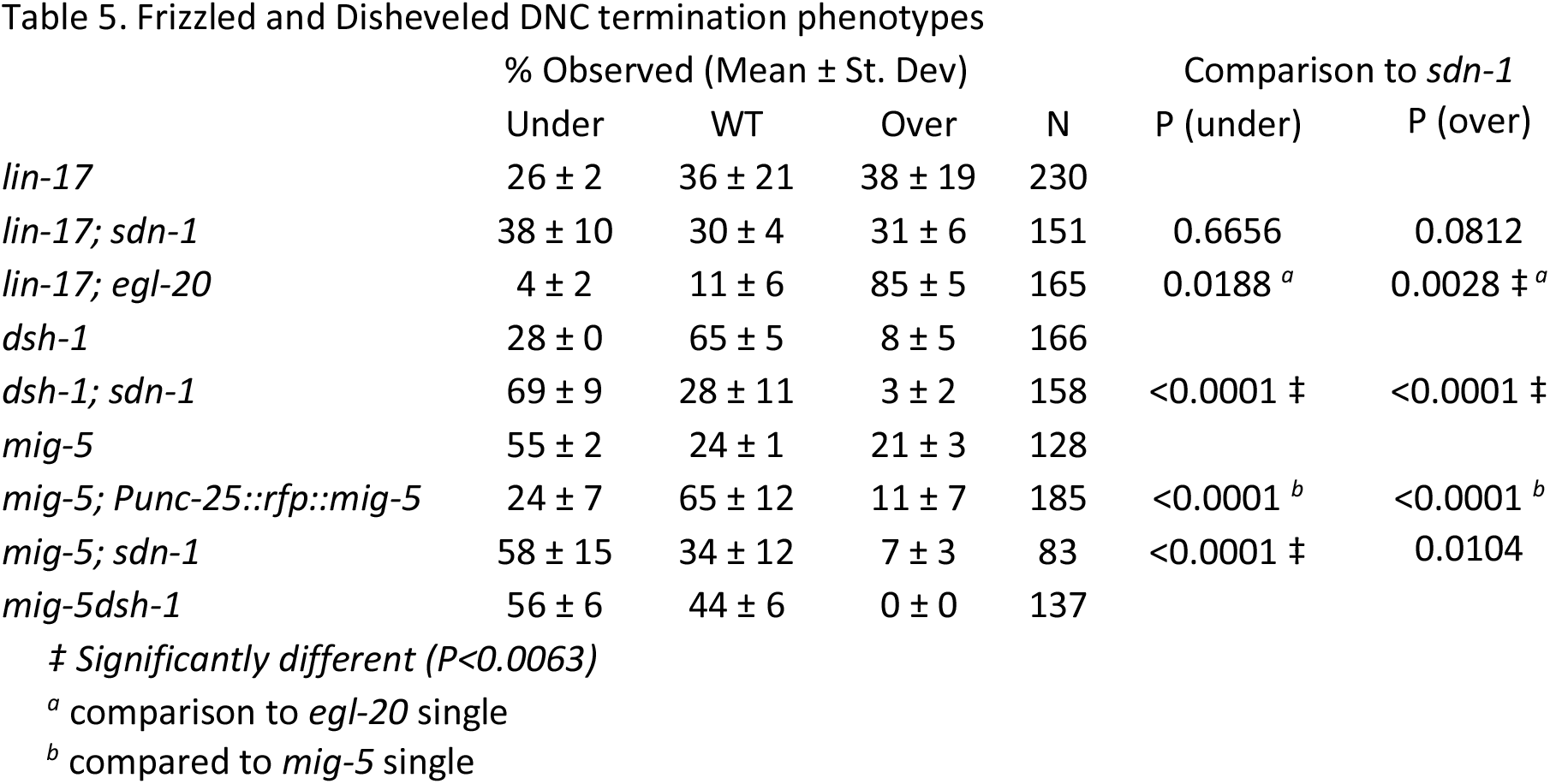
Frizzled and Disheveled DNC termination phenotypes

We created double mutants of *sdn-1* with the above Wnt signaling genes. We found that removal of *lin-17* did not grossly change the outcomes for DNC phenotypes from *sdn-1* or *lin-17* single mutants (Table 5), suggesting they were likely functioning in a common genetic pathway. Consistent with this we found that removing *egl-20* from *lin-17* resulted a reduction in the under-extension phenotype, as it had when we removed *egl-20* from *sdn-1* mutants (Table 5).

For Dishevelled, removing *sdn-1* from the *dsh-1* mutants resulted in an additive effect that was significantly different from either single mutant. The double mutants of *sdn-1* and *mig-5* were nearly synthetic lethal, with only a very small percent (~10%) of the animals surviving to adulthood. In those survivors, we observed under-extension, but it was not significantly different from *mig-5* alone. Interestingly, although *dsh-1;sdn-1* mutants were different, removing *dsh-1* from *mig-5* did not induce additional defects, as the animals appeared to be similar to *mig-5* alone. We could partially rescue the extension defects in *mig-5* animals by cell-specifically expressing *mig-5* in the GABAergic neurons, using the *unc-25* promoter (Table 5). Overall, we concluded *sdn-1* and *mig-5* function in parallel pathways during early development, but in the same pathway for DNC extension, with *mig-5* epistatic to *sdn-1.*

### LIN-17 co-localizes with SDN-1 during neurite outgrowth in an EGL-20-dependent manner

Our results suggested that LIN-17 was functioning in a manner largely equivalent to SDN-1. To determine if LIN-17 was localized to the DNC we expressed a full-length rescuing LIN-17::tagRFP chimera, under the control of the endogenous promoter (Figure 5) (Huarcaya Najarro and Ackley, 2013). LIN-17::RFP was prevalent throughout the animal, including the posterior region of the dorsal nerve cord and surrounding tissues. We examined the localization of LIN-17::RFP relative to SDN-1::GFP in the *juSi119* animals. We found an equivalent enrichment of LIN-17 and SDN-1 at the termination point of the DNC in wild-type animals.

**Figure 5.**
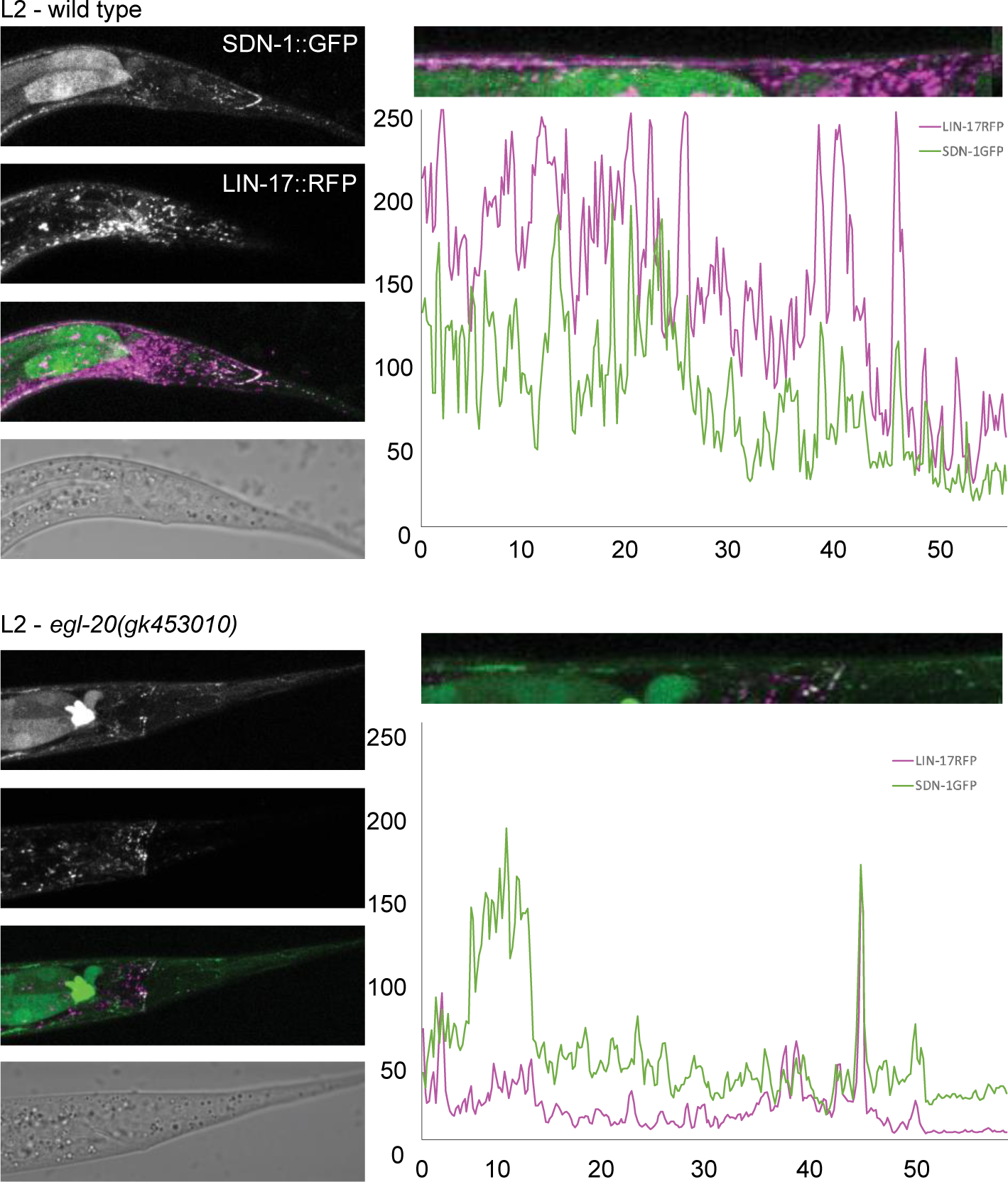
SDN-1 and LIN-17 co-localize to the dorsal nerve cord in an *egl-20-*dependent manner. Loss of function in *sdn-1* and *lin-17* had similar effects on axon outgrowth, and the proteins appear to co-localize in the dorsal nerve cord in L2 animals. Conversely, in *egl-20* loss of function animals while SDN-1::GFP localization was present, LIN-17::RFP accumulation was reduced, especially within the dorsal nerve cord.

Previous work suggested that LIN-17 localization in mechanosensory neurons was dependent on EGL-20/Wnt signaling. We asked whether this was true in the DNC by crossing the LIN-17::RFP and SDN-1::GFP transgenes into *egl-20* loss-of-function animals. We found that LIN-17 was no longer enriched in the posterior branch of the most posterior D-type neuron in young (L2) animals (Figure 5). However, later in development (L4) we could observe LIN-17 in the dorsal nerve cord in *egl-20* mutants. Thus, we concluded that, with respect to localization to the region where GABAergic axons will terminate in the DNC, LIN-17 localization was partially *egl-20-*dependent.

### SDN-1 and EGL-20 require BAR-1 function during axon outgrowth

In *C. elegans* the *bar-1*/β-catenin gene is believed to be the primary transcriptional regulator for canonical Wnt signaling (Eisenmann; Natarajan *et al.* 2001). The loss of *bar-1*/β-catenin, results in a highly penetrant under-extension phenotype (Figure 6, Table 6); (Maro *et al.* 2009)). Removing either *sdn-1* or *egl-20* from *bar-1* mutants did not change the effect of *bar-1* loss on axon termination (Table 6). Since animals with EGL-20 over-expression induced premature termination of the DNC, while loss of function mutations in *egl-20* resulted in over-extension, our results suggested EGL-20 activity likely functioned to repress BAR-1 activity in DNC outgrowth, which is not a canonical outcome of Wnt-ligand signaling. We also examined *lin-44; bar-1* double mutants and found, as had been previously described, an intermediate phenotype, with some animals having an over-extension phenotype. This is additional evidence that LIN-44 and EGL-20 were affecting the growth of the D-type neurons via distinct mechanisms.

**Figure 6.**
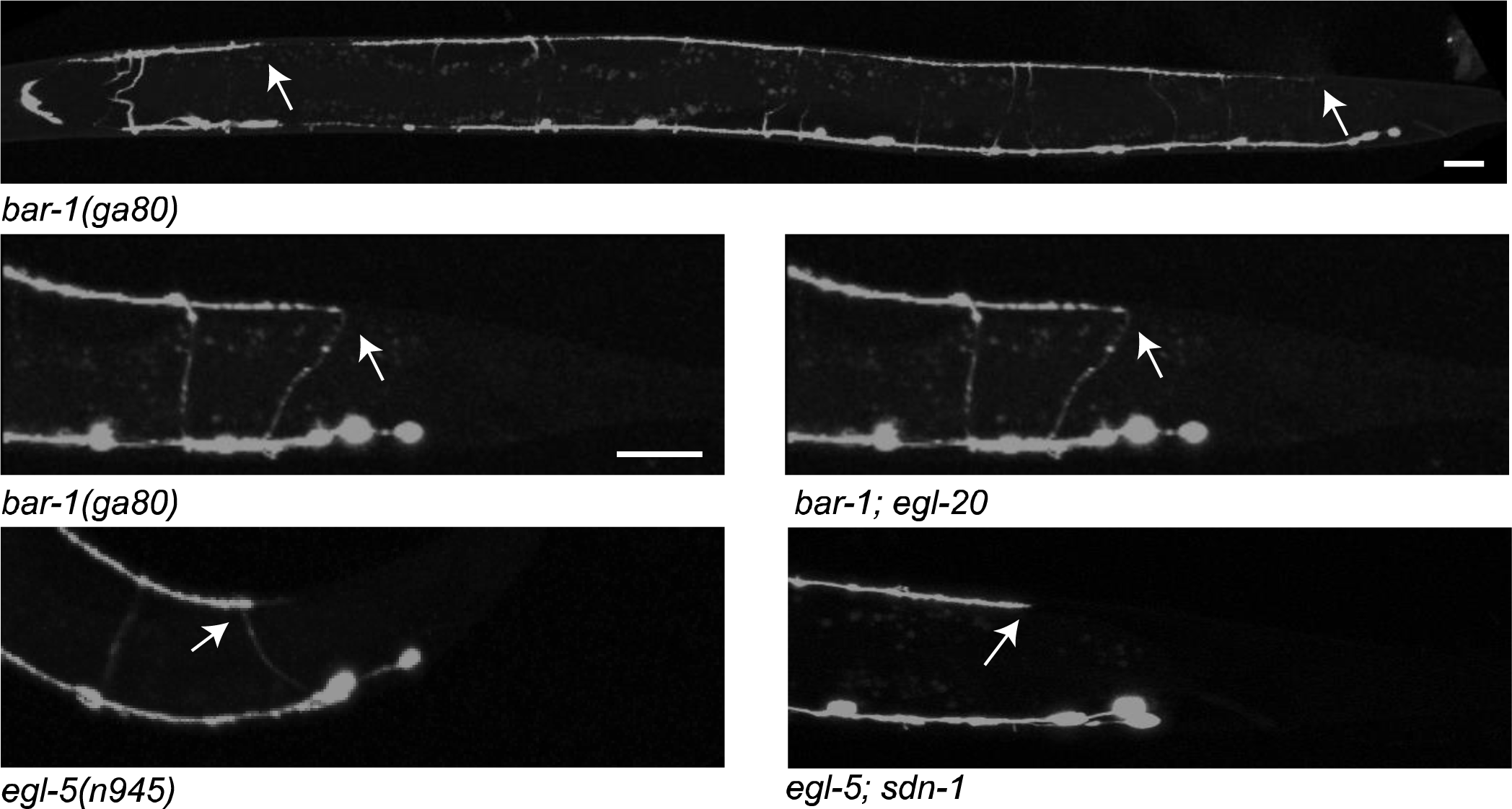
Axon underextension in *bar-1* and *egl-5* mutants. The D-type neurons in *bar-1* mutants exhibited premature termination (arrows). Specifically, in the tail, most frequently the posterior branch of the dorsal axon fails to form or was shortened. *egl-5* mutants had a similar phenotype, with an axon that grew only a short distance from the commissural branch point.

**Table 6.** *bar-1*/β-catenin DNC termination phenotypes

|  | % Observed (Mean ± St. Dev) |  |  |  | Comparison to *bar-1* |  |
| --- | --- | --- | --- | --- | --- | --- |
|  | Under | WT | Over | N | P (under) | P (over) |
| *bar-1* | 96 ± 1 | 4 ± 1 | 0 ± 0 | 196 |  |  |
| *bar-1; sdn-1* | 98 ± 2 | 2 ± 2 | 0 ± 0 | 170 | 0.5542 | 1 |
| *bar-1; egl-20* | 89 ± 4 | 10 ± 6 | 1 ± 2 | 150 | 0.0242 | 0.1901 |
| *bar-1; lin-44* | 65 ± 21 | 25 ± 19 | 11 ± 2 | 123 | <0.0001 ‡ | <0.0001 ‡ |

‡ Significantly different ( $P < 0.0125$ )

### MOM-2 is not required for initial DNC formation

Since 96% of *bar-1* animals were under-extended, we asked whether a third Wnt ligand, *mom-2*, could be responsible for promoting axon outgrowth. *mom-2* is expressed in the L1 stage, in the posterior of the animal near the region where the DNC is forming (Harterink *et al.* 2011). We examined maternally-rescued *mom-2(or77)* homozygotes and found no evidence of under-extension in the DNC, although we did see over-extension (Table 7). Removing *mom-2* did not induce under-extension in the *lin-44* or *egl-20* backgrounds. Triple mutants were sick and difficult to obtain, so we used RNAi to knockdown *mom-2* in *lin-44; egl-20* double mutants. *mom-2* RNAi caused a highly penetrant embryonic lethality, thus we collected escapers and analyzed the DNCs. We saw a higher than expected rate of under-extension in the escapers, but it was not as high as was observed in *bar-1* single mutants. Thus, it is unlikely that MOM-2 was functioning to activate BAR-1 to promote axon extension from the commissural branch point to the normal termination point. RNAi of *mom-2* in the animals lacking *sdn-1* significantly reduced the number of under-extended DNCs, similar to the effect of removing *egl-20,* which is more consistent with MOM-2 functioning to promote normal termination.

**Table 7.** *mom-2* DNC termination phenotypes

| | % Observed (Mean ± St. Dev) | | | N | Comparison to *mom-2* | | % Pdn | N |
| --- | --- | --- | --- | --- | --- | --- | --- | --- |
| *mom-2(or77) [M+]* | 0 ± 0 | 20 ± 1 | 80 ± 1 | 200 | | | 0 ± 0 | 200 |
| *mom-2(or77) [M+]; lin-44* | 0 ± 0 | 11 ± 14 | 89 ± 14 | 128 | 1 | 0.0002 | 3 ± 5 | 130 |
| *mom-2(or77) [M+]; egl-20 [M+]* | 3 ± 2 | 21 ± 25 | 76 ± 23 | 242 | 0.0344 | 0.4569 | 1 ± 2 | 243 |
| *mom-2[RNAi]* | 2 | 20 | 78 | 100 |  |  | 0 | 100 |
| *mom-2[RNAi]; lin-44; egl-20* | 16 ± 7 | 16 ± 7 | 67 ± 15 | 23 | 0.001 $\ddagger^a$ | 0.2799 <sup>a</sup> | 76 ± 2 | 95 |
| *mom-2[RNAi]; sdn-1* | 12 | 52 | 36 | 100 | <0.0001 ‡ | 0.0331 | 14 | 116 |

‡ Significantly different ( $P < 0.0063$ )
<sup>a</sup> compared to *mom-2[RNAi]*

### The EGL-5 Hox gene contributes to the initial DNC formation and outgrowth

Wnt signaling in *C. elegans* has been shown to function via many downstream mechanisms, including activating the transcription of Hox genes. The *egl-5* Hox gene is expressed in the most posterior portion of the animal, including in the posterior D-type motorneurons, and surrounding tissues (Niu *et al.* 2011) (Figure 7). A loss of function mutation in *egl-5(n945),* recapitulated the *bar-1* phenotype in the D-type neurons, causing a highly penetrant under-extended phenotype (95%). We subsequently tested double mutants of *sdn-1* with *egl-5* and found that DNCs were almost entirely under-extended (89%; P<0.001 vs. *sdn-1* and P=0.19 vs *egl-5*) (Table 8), suggesting *sdn-1* and *egl-5* functioned in the same pathway.

**Figure 7.**
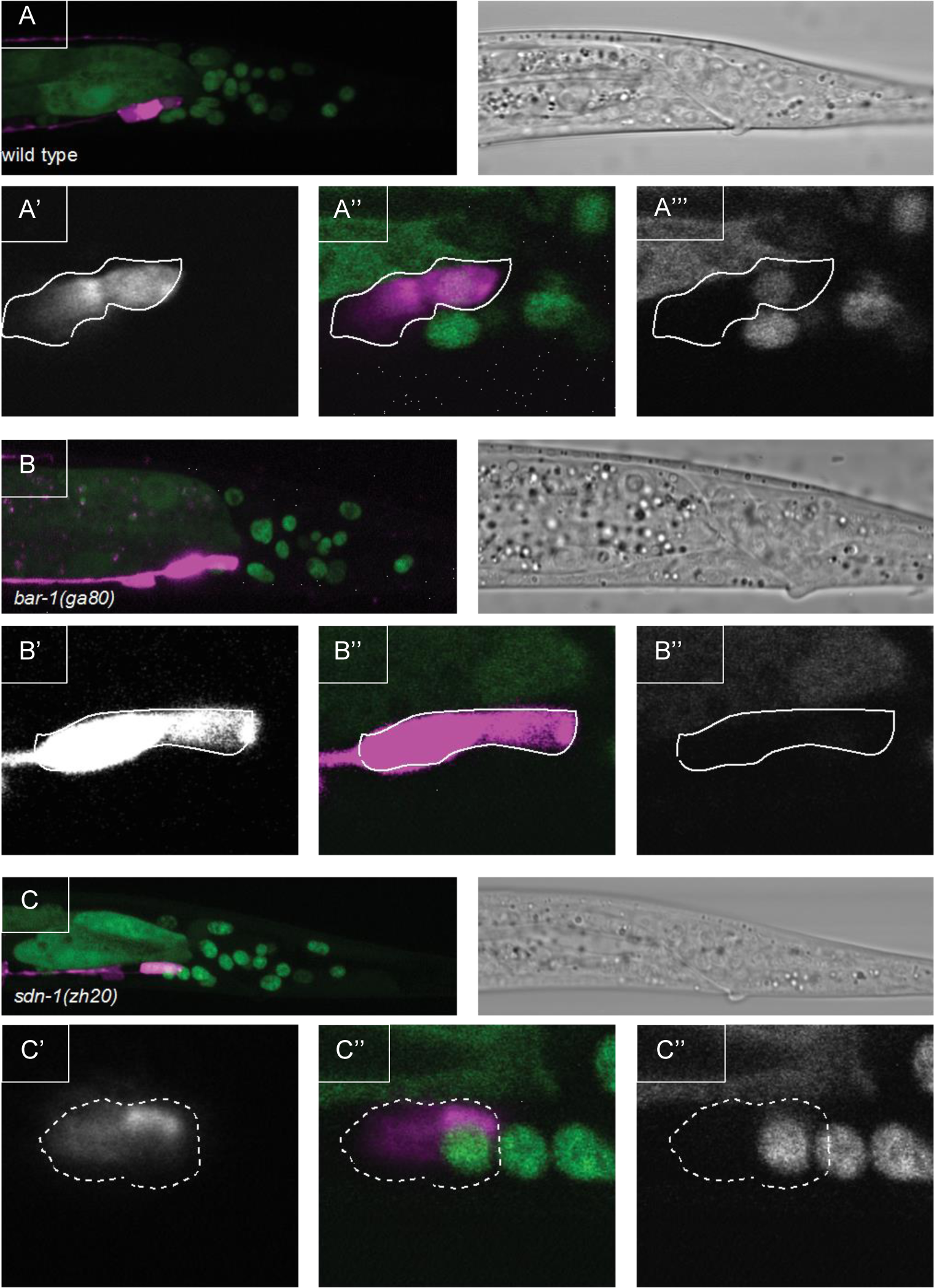
EGL-5GFP is expressed in the GABAergic motorneurons in a *bar-1*-dependent manner. We examined the expression of EGL-5 using a P*egl-5::*EGL-5::GFP transgene that rescued the mutant defects associated with the (*n945*) phenotype. A) In wild-type animals EGL-5::GFP was expressed in the most posterior D-type neurons, along with other cells in the tail. The image is a projection of a Z-stack through the tail. (A’A’) Single planes containing the VD13 nucleus are illustrated for *Punc-25::rfp* (A’) merge (A”) and *Pegl-5::egl-5::gfp* (A’). Note that EGL-5::GFP is present in the nucleus of VD13, but also in DD6 (cell anterior), but the nucleus is out of the plane capture. B) Loss-of-function in *bar-1* results in a loss of EGL-5 expression in the D-type neurons, but a complete loss of EGL-5 expression altogether. B’B’) Single plane images through the VD13 nucleus, as above. C) Loss of function in *sdn-1* does not recapitulate the *bar-1* loss of *egl-5* expression. C’-C’) Single plane images through VD13 nucleus, as above. Note, the cell bodies of DD6 and VD13 are circled in all panels.

**Table 8.**
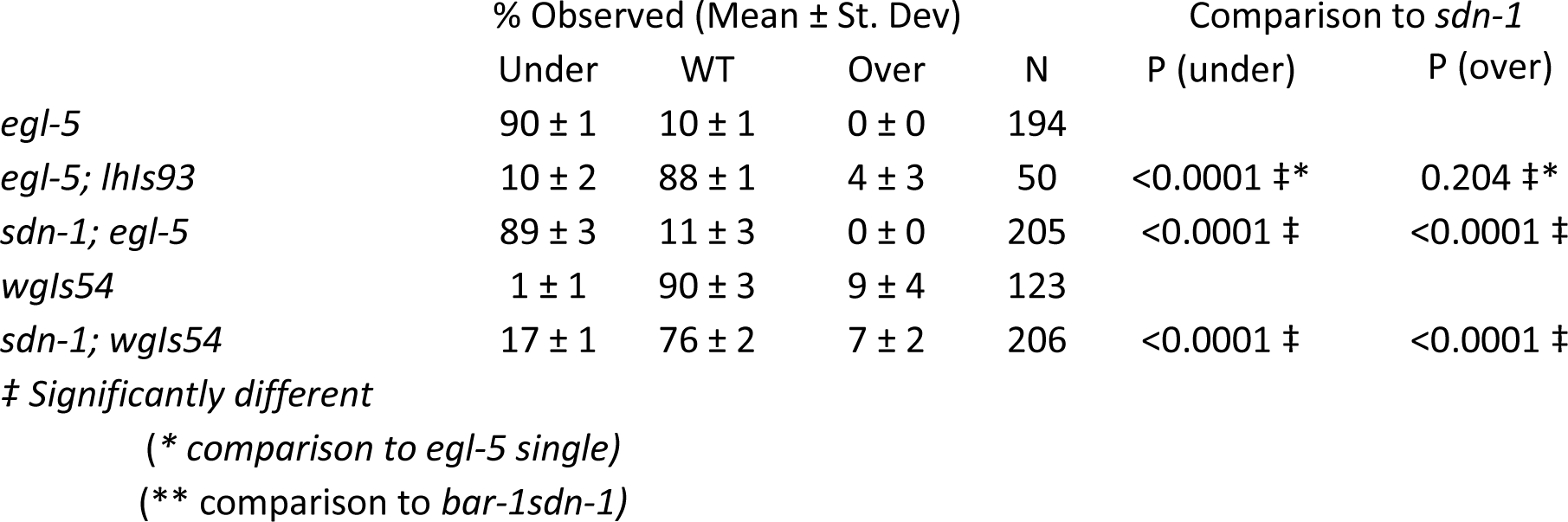
*egl-5* DNC termination phenotypes

To determine if *egl-5* was expressed in the DD and VD motorneurons we crossed an integrated P*egl-5*::EGL-5::GFP transgene (*wgIs54*) into a reporter for the GABAergic motorneurons (*Punc-25*::mCherry). We found, as has been reported, EGL-5::GFP expression was largely restricted to the posterior region. We observed co-incident expression of EGL-5::GFP and mCherry in the three most posterior neurons, VD12-DD6-VD13, with expression strongest in DD6 and VD13 (Figure 7). We observed expression in the posterior region of L1 animals, during the formation of the VD13 neuron (Figure 7), and expression was visible in the nascent VD13 (Figure 7) as soon as we could observe the differentiation marker (*Punc-25*) which occurs as the axons are initiating growth, prior to the formation of the dorsal process (Norris and Lundquist 2011) (Figure 2). Further, expression of EGL-5 was maintained in the neurons throughout development (L4 stage – Figure 7). Thus, we concluded that EGL-5 is expressed in the neurons (DD6 and VD13) that make posterior projections that terminate axon growth without the help of a more posterior D-type neuron, and that expression is maintained throughout the period of development when D-type neuron growth must be metered. We generated transgenic lines expressing *egl-5* specifically in the D-type neurons in the *egl-5(n945)* animals (*lhIs93*). Cell-specific expression could partially rescue the underextension phenotype in *egl-5* mutants (Table 8), suggesting *egl-5* function was cell-autonomous.

To better understand the genetic pathway, we crossed the *wgIs54* transgene into the *bar-1(ga80)* mutants and found that expression of the *egl-5::gfp* was lost in the DD and VD neurons, but expression could be observed in other cells adjacent (Figure 7). This suggested that EGL-5 expression in the D-type neurons was *bar-1*-dependent, and that this might underlie the under-extension defects. Similar effects of Wnt signaling activating anterior vs. posterior behaviors in cells has been documented in the migration of the Q-neuroblasts, with *mab-5* necessary and sufficient for proper posterior growth (Tamayo *et al.* 2013), and more recently in the axon outgrowth of PLM axons (Zheng *et al.* 2015a).

We subsequently examined whether *sdn-1* mutants also resulted in alterations in *egl-5* expression. In contrast to *bar-1,* we did not find EGL-5::GFP expression to be lost from the D-type neurons in *sdn-1* mutants. We did note however, the *wgIs54* transgene partially rescued axon-extension phenotypes present in the *sdn-1* loss-of-function animals (Table 8). Thus, we concluded that underextension of the DNC in *sdn-1* animals likely occurs via dysregulation of *egl-5* expression in the GABAergic neurons, and that it is possible to partially bypass the requirement for SDN-1 in early dorsal nerve cord development by increasing the levels of EGL-5. It should be noted that this observation is consistent with the differences observed in the *sdn-1* and *bar-1* mutants, in the frequency of, as well as the position where axons prematurely terminated. Both aspects were more severe in *bar-1* mutants than in *sdn-1*, and thus we do not believe *sdn-1* mutations resulted in a complete loss of either *bar-1* or *egl-5* activity.

## Discussion

### SDN-1 functions in anterior-posterior axon outgrowth and termination

Neurons form neural networks by extending axons and dendrites into discrete regions where they can form synapses with targets. The complexity of network formation relies on extracellular cues being interpreted by cell-surface receptors, often with great precision and fidelity. Here we have found that two neuron types that terminate axon growth at stereotyped positions in the *C. elegans* nervous system are affected by the loss of the cell-surface receptor Syndecan/SDN-1. These results are consistent with *sdn-1* regulating cell migration along the A/P axis (Rhiner *et al.* 2005; Schwabiuk *et al.* 2009). The disparate termination points and function of the distal tip cells, the GABAergic motorneurons and the posterior mechanosensory neurons, suggest that SDN-1 provides migration termination information broadly during organismal development.

In both GABAergic and mechanosensory neurons we found axons stopping short of, and growing past, their stereotyped termination point, even when other aspects (initial outgrowth, dorsal/ventral guidance, *etc.*) were grossly normal. Therefore, we concluded that the termination defects were, at least partially, independent of earlier guidance errors. Neither the PLMs nor the terminal D-type neurons (DD6 and VD13) have obvious cellular landmarks that indicate where they should terminate. This contrasts with the more anterior DD and VD neurons that terminate their posterior dorsal branch on the next DD or VD cell posterior, that is, the posterior branch of the VD11 dorsal process terminates where it meets the anterior branch of VD12, *etc*. Because of the overlap of the DD and VD axons in the dorsal nerve cord, it is not simple to determine whether *sdn-1* has a broader role in regulating the outgrowth of dorsal branches in the D-type neurons along the anterior-posterior axis of the animal, and thus we focused solely in this work on those in the posterior-most region. Our results suggest that SDN-1 modulates Wnt signaling to facilitate axon outgrowth early in development, and subsequently to maintain axon extension to be allometric with organismal growth as development progresses.

### SDN-1 function is both autonomous and non-autonomous

In the D-type neurons we found an early role for SDN-1 in ensuring that the posterior neurite of the dorsal nerve cord reached the proper termination point. Based on rescue experiments, SDN-1 appeared to function cell-nonautonomously, as we could rescue the defects by expressing SDN-1 with an *sdn-1* promoter, but not when we replaced SDN-1 specifically in the D-type neurons (Figure 8). We found that *sdn-1* mutants exhibited an over-growth phenotype, where the axons grew past the normal termination point. This phenotype appeared to be cell-autonomous, as we could largely rescue this phenotype by specifically replacing SDN-1 in the GABAergic neurons. Thus, we concluded that SDN-1 is necessary in the D-type neurons to prevent overgrowth. Within the D-type neurons the PDZ-binding motif (EFYA) was required for the proper localization and function of SDN-1.

**Figure 8.**
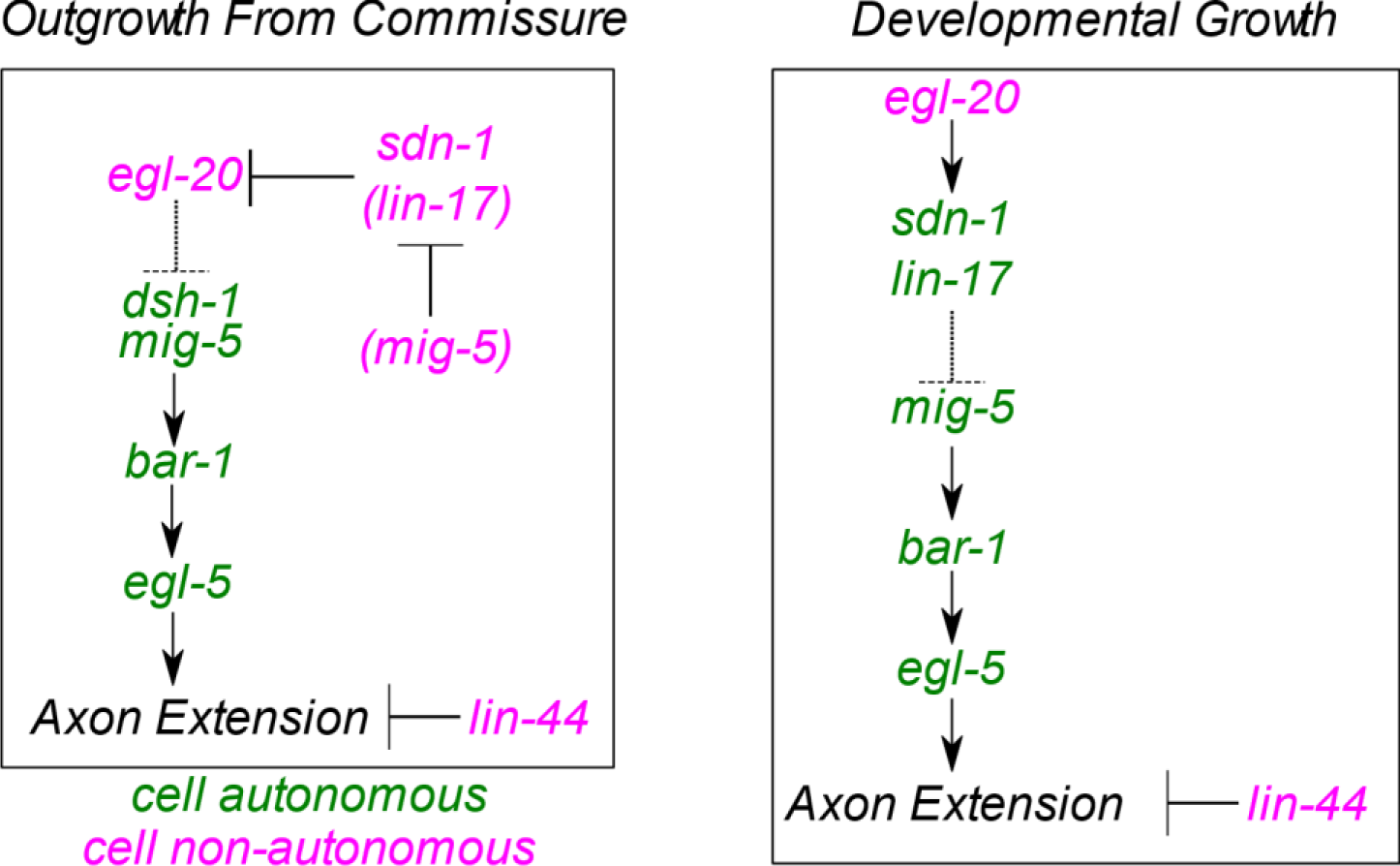
Genetic model. A simplified genetic model suggests that *egl-20* and *lin-44* exhibit independent effects on the extension of axons into the posterior of the animal. In the *egl-20* pathway, we found a cell-nonautonomous role for *sdn-1* in the early outgrowth of the commissure, where *sdn-1* appears to inhibit the activity of *egl-20.* We infer that *mig-5* and *lin-17* have non-autonomous roles, due to the phenotypic consequences and the lack of complete rescue for *mig-5* by cell-specific expression. Within the D-neurons the link between *egl-20* appears to inhibit the activities of *dsh-1, mig-5, bar-1* and *egl-5,* all of which promote axon extension. Later in development, as the animal is increasing in size, the function for *sdn-1* appears cell autonomous, and it now appears to function with *egl-20* to inhibit axon extension, by inhibiting the activity of the *mig-5, bar-1, egl-5* components that promote outgrowth.

One result that was somewhat curious was the rescue of *sdn-1* phenotypes by the integrated transgenes with the endogenous *sdn-1* promoter. We could rescue the under-growth phenotype, and other *sdn-1* phenotypes, including uncoordinated movement and male mating (data not shown), but not the over-extension phenotype. We found no evidence that SDN-1 expressed from the integrated transgenes was expressed in the D-type neurons, despite previous reports that the gene is expressed there (Rhiner *et al.* 2005). It is certainly possible that the levels of expression are below detection, or that expression was delayed relative to when we conducted our analyses. However, we did find that we could rescue the over-extension by D-type cell-specific expression. Thus, we concluded that, for reasons that are not yet understood, two, separately developed SDN-1 MosSCI inserted transgenes, were not being well expressed in the D-type neurons. Other transgenes integrated via MosSCI at that position do express well in the D-type neurons (BDA, personal observations), and thus it is not clear why the SDN-1 transgenes are not.

### SDN-1 functions with EGL-20 to regulate axon termination

Our *sdn-1* rescue experiments did confirm that both the under- and over-growth phenotypes were due to loss of function in *sdn-1.* Further, the cell-specific expression of *sdn-1* suggests a model whereby SDN-1 is acting outside the D-type neurons to facilitate growth from the commissural branch point to the normal termination point. In animals lacking both *sdn-1* and *egl-20* we found fewer animals with under-extended axons, suggesting that SDN-1 was negatively regulating EGL-20 during the initial axon outgrowth. This is consistent with our evidence and previous reports that EGL-20 primarily functions to inhibit axon outgrowth. Support for this hypothesis came from our observation that we could induce premature axon termination by over-expressing EGL-20, and that we could exacerbate the effects of EGL-20 over-expression by removing SDN-1, consistent with an antagonistic role for SDN-1 in this event.

We found that SDN-1 appeared to be aggregated near the site where axons would initially terminate. A simple model then would be that SDN-1 regulates the local accumulation of EGL-20 near the presumptive axon termination point. However, while the axon terminates near places we see EGL-20 accumulating, we do not necessarily see EGL-20 becoming enriched near the termination point, relative to adjacent regions. Thus, it is possible that SDN-1 accumulation has additional functions, either by regulating the accumulation of other ligands, or forming a complex with EGL-20 that has differential activity with respect to axon termination. regulates the accumulation of multiple, coordinated ligands at the termination point, and it is the aggregate activity of these ligands that terminates axon growth.

After the initial role in antagonizing EGL-20, we found that SDN-1 was required cell-autonomously to promote termination of the axon during allometric growth of the D-type neurons. In this capacity, SDN-1 and EGL-20 appear to function equivalently, as loss of function in either resulted in axons that could grow past the normal termination point. Our epistasis experiments suggested that *sdn-1* acted genetically downstream of EGL-20/Wnt, but with the LIN-17/Frizzled, and upstream of the MIG-5/Dishevelled, BAR-1/β-catenin and EGL-5/Hox transcription factor. Similar types of results suggesting a co-receptor role that can both promote and inhibit Wnt signaling have previously been demonstrated for the CAM-1/Ror tyrosine kinase, which has been referred to as a Wnt co-receptor (Forrester *et al.* 1999; Forrester *et al.* 2004; Green *et al.* 2007; Hayashi *et al.* 2009; Kennerdell *et al.* 2009; Ohama and Hayashi 2009; Minami *et al.* 2010; Song *et al.* 2010). Interestingly, the cell-autonomous function of SDN-1 required the EFYA motif, which, in vertebrates, links syndecans to PDZ-domain containing proteins. Here we found that MIG-5/Dishevelled is required for axon outgrowth. MIG-5, like most Dishevelled proteins, contains a PDZ domain, central to the protein. Whether MIG-5 and SDN-1 physically interact remains to be determined empirically, but if they do, our evidence would suggest SDN-1 acts to inhibit MIG-5 to limit axon outgrowth.

### HS modifiers regulate the activity of EGL-20

SDN-1 is the most prominent HSPG in *C. elegans* as detected by an antibody that recognizes the stubs of HS side chains resulting from heparitinase treatment (Hudson *et al.* 2006). We found that enzymes that modify the heparan sulfate side chains, partially recapitulated the loss of function in *sdn-1,* but likely represent the loss of HS-chains on multiple core proteins, as *sdn-1* and *hse-5* double mutants had an additive effect. However, we did not find evidence that those core proteins are the glypican-like receptors, suggesting perhaps another HSPG is functioning in this pathway. With the primary candidates for this being UNC-52/perlecan and CLE-1/collagen XVIII.

Like the loss of *sdn-1*, removing *egl-20* from animals with mutations in the modifier enzymes resulted in fewer under-extended DNCs, consistent with an apparent increase in EGL-20 activity (or range of activity) when animals had deficits in HS-modification. Our results are consistent with the modifications being important in mediating proteoglycan interactions with EGL-20. Interestingly, EGL-20 has a poly-basic region (AA288-303 RKATKRLRRKERTERK) that is absent from the other posteriorly expressed Wnt ligands LIN-44 and MOM-2. Thus, it is conceivable that EGL-20 interactions are electrostatically favorable with proteins that have been modified with acidic-HS side chains, although this would require a more thorough biochemical analysis and remains hypothetical at this point.

### Wnt ligands antagonize but BAR-1 and EGL-5 promote posterior axon outgrowth

In canonical signaling the Wnt ligand binding to Frizzled results in increased β-catenin activity. Here, although *egl-20* appears to inhibit growth, we find that *bar-1*/β-catenin activity promotes axon outgrowth. Interestingly, in the *bar-1* mutants, the posterior branch of the axon in the dorsal cord is almost entirely absent, although the anterior branch is typically intact. This suggests that perhaps, *bar-1* is required for the earliest events of axon outgrowth in the posterior branch, although how that would be differentially regulated from the formation of the anterior branch is curious. Ultimately, our results are consistent with previous work demonstrating that *bar-1* acts as a switch for axon growth. When *bar-1* is absent axons are shortened, and when *bar-1* degradation is inhibited axons are overgrown (Maro *et al.* 2009).

Our results suggested that Wnt ligands were not functioning to activate BAR-1 signaling, rather, it appears to be the opposite. In the PLM neurons, recent work has demonstrated that Dishevelled proteins likely regulate the repellant activities of Wnt signaling, by modulating the phosphorylation state of Frizzleds (Zheng *et al.* 2015a). However, we find that, in the D-neurons, there are some important differences, including opposite effects of loss of Wnt ligands and Disheveled. Further, in the PLM neurons there was only a modest effect on axon growth due to the loss of *bar-1,* but we found a strong effect on GABAergic outgrowth. One possibility is that the effects are somehow different on the anteriorly growing PLM axons, compared to the posteriorly directed D-type axons.

Thus, it is not yet clear how BAR-1 is activated in the D-type neurons to promote axon outgrowth, nor how Wnt signaling is inhibiting BAR-1 signaling in these neurons. *rpm-1*(an E3 ubiquitin ligase) and *anc-1*(a Nesprin family member) also appear to function in neurons to potentiate *bar-1* signaling (Tulgren *et al.* 2014). Thus, it is possible that, some amount of Wnt ligand activity initiates BAR-1 signaling, and this can support axonal development due to the function of RPM-1 and ANC-1.

In the D-type neurons BAR-1 appeared to be necessary to activate the expression of EGL-5, and loss of *egl-5* activity largely recapitulates the *bar-1* mutant phenotype. Other Hox genes, for example *mab-5*, have been shown to be both necessary and sufficient for the posterior growth of the Q neuroblast descendants (Tamayo *et al.* 2013). Thus, it is possible that *egl-5* has a similar role in the D-type neurons. However, we did not necessarily observe increased axon outgrowth in the anterior of the animal when EGL-5 is driven ectopically in the anterior D-type neurons. This could be due to a different program of axon outgrowth in those neurons, or that EGL-5 is more specifically required for axon outgrowth in the posterior D-type neurons, but that it is most easily observed in the posterior branch of the dorsal cord.

### Wnt-dependent mechanism of axon termination

The wire-minimization hypothesis suggests that axon pathfinding has evolved to provide a balance between the length of process, which reduce neuronal efficiency, and the necessary routing (Miller-Fleming *et al.* 2016). Consequently, it is likely that organisms evolved multiple, redundant signals to regulate aspects of pathfinding, including where axons will choose to terminate. Wnt signaling is a primary force in the anterior-posterior guidance of neurons across the animal kingdom (Zou 2004; Fenstermaker *et al.* 2010). Here we have found that multiple Wnt ligands collaborate with the syndecan HSPG to regulate the termination of D-type neurons in a stereotyped position. Although these molecules demonstrate a somewhat complicated pattern of genetic interactions, our results are consistent with them being necessary to reproduce the stereotyped termination of the GABAergic motorneurons in the dorsal nerve cord.

## Acknowledgements

We would like to thank Katsu Dejima and Andrew Chisholm for sharing the *juSi119* MosSCI insertion prior to publication. We would also like to thank Riley Roberts, Vi Leitenberger, Erik Lundquist and Michael Branden for useful discussion and technical support during this work. This work was supported in part by awards from the NIH (P20GM103418 and P20GM103638).

